# Using *Plasmodium knowlesi* as a model for screening *Plasmodium vivax* blood-stage malaria vaccine targets reveals new candidates

**DOI:** 10.1101/2020.08.07.241125

**Authors:** Duncan N. Ndegwa, Jessica B. Hostetler, Alejandro Marin-Menendez, Theo Sanderson, Kioko Mwikali, Lisa H. Verzier, Rachael Coyle, Sophie Adjalley, Julian C. Rayner

## Abstract

*Plasmodium vivax* is responsible for the majority of malaria cases outside Africa. Unlike *P. falciparum*, the *P. vivax* life-cycle includes a dormant liver stage, the hypnozoite, which can cause infection in the absence of mosquito transmission. An effective vaccine against *P. vivax* blood stages would limit symptoms and pathology from such recurrent infections, and therefore could play a critical role in the control of this species. Vaccine development in *P. vivax*, however, lags considerably behind *P. falciparum*, which has many identified targets with several having transitioned to Phase II testing. By contrast only one *P. vivax* blood-stage vaccine candidate based on the Duffy Binding Protein (PvDBP), has reached Phase Ia, in large part because the lack of a continuous *in vitro* culture system for *P. vivax* limits systematic screening of new candidates. We used the close phylogenetic relationship between *P. vivax* and *P. knowlesi*, for which an *in vitro* culture system in human erythrocytes exists, to test the scalability of systematic reverse vaccinology to identify and prioritise *P. vivax* blood-stage targets. A panel of *P. vivax* proteins predicted to function in erythrocyte invasion were expressed as full-length recombinant ectodomains in a mammalian expression system. Eight of these antigens were used to generate polyclonal antibodies, which were screened for their ability to recognize orthologous proteins in *P. knowlesi*. These antibodies were then tested for inhibition of growth and invasion of both wild type *P. knowlesi* and chimeric *P. knowlesi* lines modified using CRISPR/Cas9 to exchange *P. knowlesi* genes with their *P. vivax* orthologues. Candidates that induced antibodies that inhibited invasion to a similar level as PvDBP were identified, confirming the utility of *P. knowlesi* as a model for *P. vivax* vaccine development and prioritizing antigens for further follow up.

**AUTHOR SUMMARY:** Malaria parasites cause disease after invading human red blood cells, implying that a vaccine that interrupts this process could play a significant role in malaria control. Multiple *Plasmodium* parasite species can cause malaria in humans, and most malaria outside Africa is caused by *Plasmodium vivax*. There is currently no effective vaccine against the blood stage of any malaria parasite, and progress in *P. vivax* vaccine development has been particularly hampered because this parasite species cannot be cultured for prolonged periods of time in the lab. We explored whether a related species, *P. knowlesi*, which can be propagated in human red blood cells *in vitro*, can be used to screen for potential *P. vivax* vaccine targets. We raised antibodies against selected *P. vivax* proteins and testedtheir ability to recognize and prevent *P. knowlesi* parasites from invading human red blood cells, thereby identifying multiple novel vaccine candidates.

## INTRODUCTION

Malaria remains a major global health challenge, with an estimated 228 million cases and >400,000 deaths in 2018 (1). While there are five *Plasmodium* species that can cause malaria in humans, the majority of clinical cases are caused by *P. falciparum* and *P. vivax. P. falciparum* causes almost all malaria cases in Africa, but *P. vivax* is the dominant cause of malaria in the Americas, and causes a similar number of cases as *P. falciparum* in South-east Asia (1)⍰. As well as having different global distributions, the two species are also very different biologically, which has significant implications for control. *P. vivax*, along with *P. ovale*, can form hypnozoites during its liver stage, which are quiescent forms of the parasite that remain dormant from weeks to years in the liver, re-emerging upon stimulation to cause a relapse of malaria. Hypnozoites can therefore act as a continuous source of infection even in the absence of active transmission. This hurdle is made more significant by the fact that primaquine and tafenoquine, the only drugs used to treat hypnozoites, are frequently contraindicated due to their toxicity in patients with glucose-6-phosphate deficiency, a common polymorphism in regions of the world where *P. vivax* is most prevalent (2,3)⍰. In addition, sexual stage development in *P. vivax* is much more rapid than in *P. falciparum (4,5),* meaning that even with rapid treatment with antimalarials, onwards transmission can still occur. These features limit the effectiveness of current chemotherapeutic interventions, making the search for an effective vaccine even more important for *P. vivax*.

The complex life-cycle of *Plasmodium vivax* parasites present multiple potential intervention strategies, including preventing transmission to the mosquito, targeting the liver stage to prevent disease and relapse, and targeting blood stages to limit disease and lower the potential of transmission from one infected individual to another. Indeed, vaccine targets across all these various stages of the parasite are under investigation, although in general far fewer antigens have been studied in depth in *P. vivax* relative to *P. falciparum* (reviewed in (6,7)). This is particularly the case for blood stage targets, where only a few targets such as *P. vivax* apical membrane antigen 1 (PvAMA1) (8,9)⍰ and *P. vivax* merozoite surface protein 1 (PvMSP1_19_) (10,11)⍰ have advanced to pre-clinical study. The furthest advanced *P. vivax* vaccines, by far, are based on *P. vivax* Duffy Binding Protein (PvDBP), the only blood stage target that has reached clinical Phase Ia trials (12); these are PvDBPII-DEKnull (12)⍰ and PvDBPII (11–15)⍰⍰⍰. This is in stark contrast to *P. falciparum*, where multiple targets in different stages have been tested in Phase Ia (Reviewed in (6)⍰), and the RTS,S pre-erythrocytic vaccine has advanced beyond Phase III to pilot testing across three countries in Africa (16). More *P. vivax* targets clearly need to be screened if vaccine development for this species is to advance.

It was previously believed that *P. vivax was* completely dependent on the interaction between PvDBP and its receptor, Duffy Antigen Receptor for Chemokine (DARC) (17–20) to invade human erythrocytes. However, it has recently been shown that *P. vivax* is also able to infect individuals who are Duffy negative, so express little or no DARC on the surface of their erythrocytes (21–24). While the invasion of Duffy negative erythrocytes could still rely on PvDBP (25,26), the sole focus on PvDBP as a vaccine candidate clearly needs to be reassessed and additional targets evaluated, either as potential substitutes for PvDBP, or to be used in combination with it. As noted above, erythrocyte invasion is a very complex process, and while the process is much less well-understood in *P. vivax* than it is in *P. falciparum* (27)⍰, other *P. vivax* ligands such as reticulocyte-binding protein 2 (RBP2b) (28), GPI-anchored micronemal antigen (GAMA) (29)⍰, and erythrocyte binding protein 2 (ebp2) (30) have all been shown or proposed to be involved, enabling the identification of possible combinatorial vaccines (31)⍰.

In this study we took a reverse vaccinology approach to identify new *P. vivax* vaccine targets, building on previous work where we expressed a panel of 37 full-length recombinant *P. vivax* vaccine targets predicted to be involved in erythrocyte invasion (32)⍰. Polyclonal antibodies were generated against 8 of these proteins, and the antibodies were tested for their ability to inhibit merozoite invasion. *P. vivax* preferentially invades immature erythrocytes (33)⍰ which are difficult to obtain (34–37), which has limited the development of continuous culture of *P. vivax in vitro*, despite herculean efforts (38). As a first-stage screen we therefore performed invasion inhibition assays using *P. knowlesi*, a close phylogenetic relative of *P. vivax* (39,40) that has been adapted to *in vitro* cell culture in human erythrocytes (41,42). We also took advantage of the amenability of *P. knowlesi* to genetic manipulation to explore the function of some of the target genes, and to swap *P. knowlesi* genes for their *P. vivax* orthologues to establish whether this would affect antibody inhibition. Together, this work prioritises new targets for *P. vivax* vaccine development, and presents additional evidence that *P. knowlesi* can be used as a readily manipulatable *in vitro* model for *P. vivax*.

## RESULTS

### Generation of polyclonal antibodies against new *P. vivax* vaccine candidates

We have previously expressed a pilot library of 37 *P. vivax* proteins that were either shown to localise to merozoite organelles with a role in invasion, or were predicted to do so based on the localisation of their respective *P. falciparum* homologues (32). In all cases, the full-length extracellular domains of these proteins were expressed using a mammalian protein expression system. This approach, which increases the likelihood of correct folding of disulphide-linked extracellular domains, has been used extensively for *P. falciparum* invasion-associated proteins (43) to generate antigens capable of inducing potent invasion-inhibitory antibodies, including for the major *P. falciparum* blood-stage vaccine target PfRh5 (31,44)⍰. Comparing immunoreactivity of native or heat-denatured epitopes and testing for protein-protein interactions indicated that the produced *P. vivax* library was also likely to consist of largely functional proteins (32)⍰. To test whether this library could also be used to generate inhibitory antibodies, rabbit polyclonal antibodies were raised against eight targets, selected to represent a range of predicted subcellular localizations and including PvDBP as a positive control (Table 1). In all cases, antibodies were raised against the complete recombinant ectodomain of the candidates. As outlined in the Methods, 1mg of antigen was used to immunize each rabbit, and total IgG purified from serum using Protein A affinity chromatography.

### Anti-*Plasmodium vivax* antibodies are able to recognise orthologues in *P. knowlesi*

Given the inherent difficulty in acquiring *P. vivax ex vivo* isolates for testing, we explored the use of *P. knowlesi, which has recently been adapted to continuous in vitro cell culture in human erythrocytes (41,42) as a model for P. vivax vaccine candidate screening. P. knowlesi*, which falls into the same clade of simian parasites as *P. vivax* (39,40) naturally infects the Kra cynomolgus macaque (*Macaca fascicularis*) but causes severe zoonotic malaria in Southeast Asia (45), falls into the same clade of simian parasites as *P. vivax* (39,40). While this phylogenetic relationship is reflected in a higher of conservation between the *P. vivax* and *P. knowlesi* genomes than the *P. vivax* and *P. falciparum* genomes (39), the degree of conservation varies between at the individual gene level. Sequence alignment between our *P. vivax* candidate proteins and their orthologues in *P. knowlesi* showed a range of sequence similarities (Table 1), from a pairwise identity score of 51% for PvDBP and its closest *P. knowlesi* orthologue PkDBPα, to higher identity scores for several targets (PvGAMA, Pv12, PvARP, PvCyRPA and Pv41), reaching 80% in the case of Pv41. In contrast, a lower degree of conservation was found for the merozoite surface proteins PvMSP3.10 and PvMSP7.1, both members of multigene families which are known to be highly polymorphic within and between *Plasmodium* species.

To explore whether variable degrees of homology would limit our ability to test specific targets in *P. knowlesi*, we first determined whether antibodies raised against *P. vivax* (Pv) targets can recognise their *P. knowlesi* (Pk) orthologues in immunoblots using *P. knowlesi* schizont-stage protein lysates (Figure 1).

**Figure 1.**
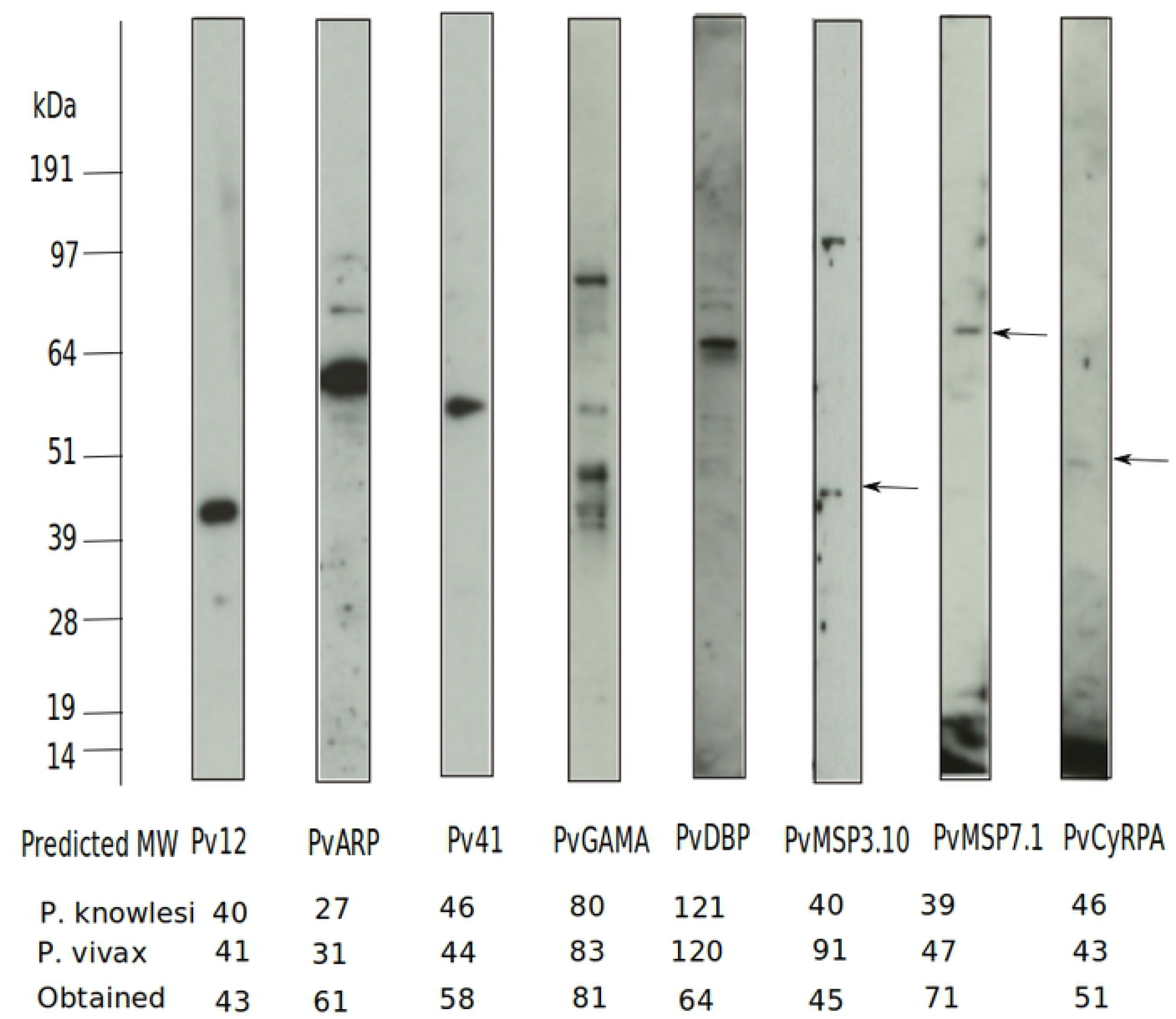
Anti-*P. vivax* polyclonal antibodies recognise proteins in *P. knowlesi* schizont protein lysates. Protein extracts from enriched *P. knowlesi* cultures at schizont stage were blotted with various anti-P. Vivax polyclonal antibodies individually and detected using ECL prime Western Blotting detection reagent, (GE Healthcare). On the left is a molecular marker in kilodalton (Kda). Obtained size (the lower row) correspond to the major bands seen. This is compared to the expected size of the corresponding protein in *P. knowlesi (upper row) and its orthologue in P. vivax* (middle row). The arrows indicate the major bands for PvMSP3.10, PVMSP7.1 and PvCyRPA. Note that the expected size of PkMSP7.1 of 39Kda is an average of the molecular weight of four PkMSP7 like proteins (i.e. PKNH_1265900, PKNH_1266000, PKNH_1266100, PKNH_1266300) that range from 32 - 46 Kda.

Antibodies raised against Pv12, PvARP, Pv41, PvMSP7.1 and PvDBP produced a single immuno-reactive band, while several bands were detected with antibodies against PvGAMA, PvMSP3.10 and PvCyRPA, suggesting either post-translational modifications or proteolytic processing events. While multiple factors could affect signal strength, including expression level in schizont stages, there was a correlation between % Pv/Pk identity and the strength of the immunoblot signal, PvCyRPA being the exception with a weak detection signal despite 68% identity. Antibodies against Pv12, PvGAMA, PvMSP3.10, PvCyRPA all detected proteins around their expected molecular weight based on estimates from the corresponding orthologous protein in *P. knowlesi*. In contrast, anti-PvARP, Pv41 and PvMSP7.1 detected proteins larger than the expected molecular weight suggesting that they might migrate more slowly, which is not uncommon in extracellular proteins. Anti-PvDBP detected a protein half the expected size suggesting either that PkDBPα (the closest *P. knowlesi* orthologue of PvDBP) is highly processed, or that anti-PvDBP antibodies cross-react with multiple PkDBP proteins. Overall however, immunoblotting showed that the majority of anti-*P. vivax* antibodies recognised *P. knowlesi* proteins.

### Anti-*P. vivax* antibodies are able to localise orthologous target proteins to *P. knowlesi* invasion organelles

To further explore the use *of P. knowlesi* as a model for *P. vivax* reverse vaccinology studies, we tested the antibodies in indirect immunofluorescence assays using mature *P. knowlesi* schizonts (Figure 2). Out of the 8 polyclonals, only anti-PvMSP3.10 (which shows the lowest percentage of identity between Pv and Pk) did not produce a specific signal (Figure 2). Anti-PvGAMA, PvCyRPA, PvDBP and PvARP all labelled punctate foci within the merozoites, while anti-Pv12, Pv41 and PvMSP7.1 all appeared to label the entire merozoite surface. No staining was observed with pre-immune antiserum (Figure S1), confirming that the labelling was antigen-specific.

**Figure 2.**
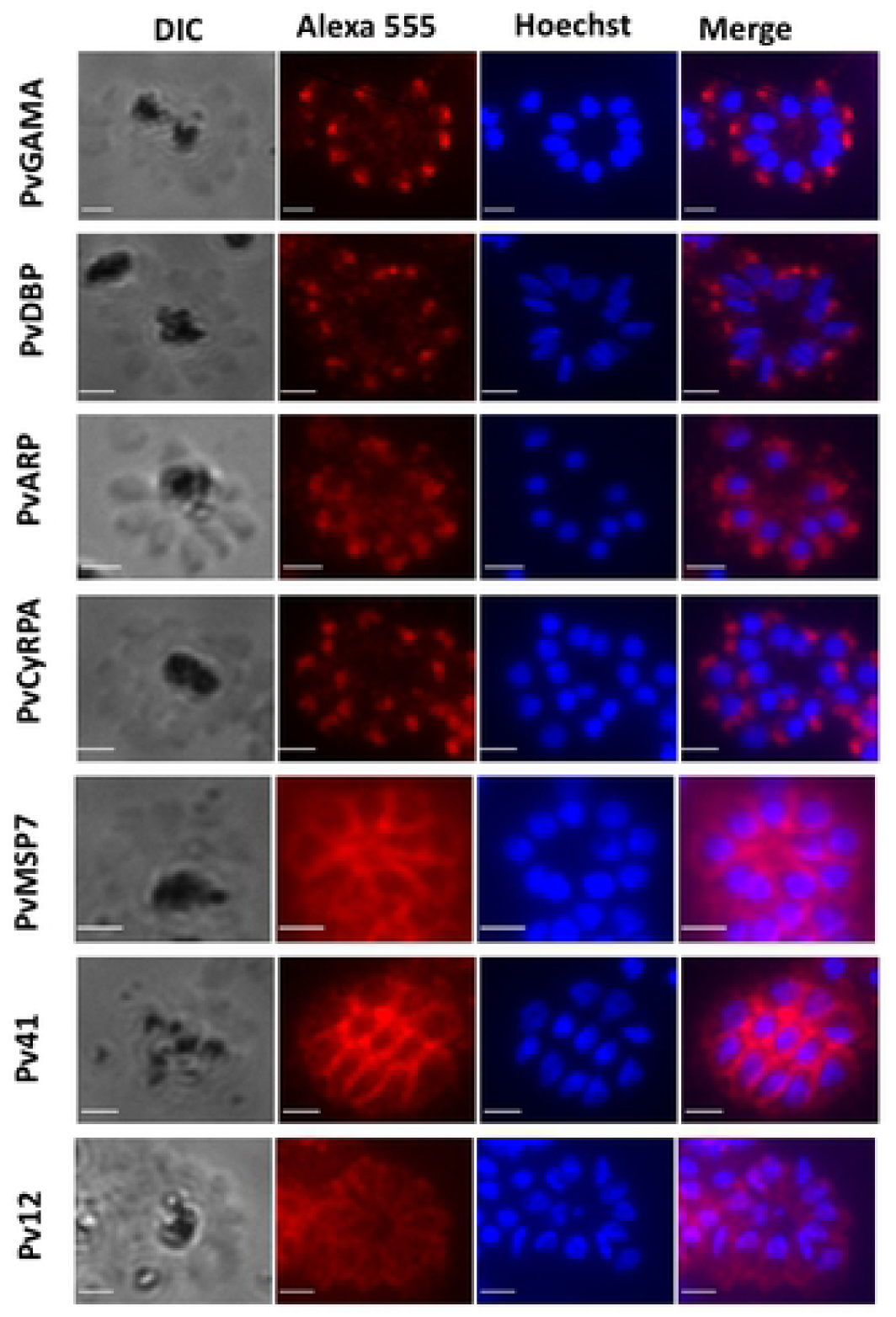
Immunolocalisation of *P. knowlesi* proteins using polyclonal anti-*P. vivax* antibodies. *P. knowlesi* homologs of *P. vivax* vaccine candidates were localised using *P. vivax* polyclonal antibodies and Alexa Fluor 555 Goat-anti rabbit (Thermo Fisher Scientific) as the secondary antibody. Localisation of PkGAMA, PkDBP, PkARP, PkCyRPA, PkMSP7.1, Pkp41 and Pkp12 was performed using antibodies against PvGAMA, PvDBP, PvARP, PvCyRPA, PvMSP7.1, Pv41 and Pv12, respectively. Parasite nuclei were stained with Hoechst 33342 (Thermo Fisher Scientific). Merge is an overlay of Alexa 555 and Hoeschst. Scale bar is 2 micrometers.

To establish the specific location of each antigen, anti-*P. vivax* antibodies were used in co-localization experiments with antibodies specific to proteins of known cellular locations i.e. AMA1, MSP1-19 and GAP45 located in microneme, merozoite surface and inner membrane complex (IMC), respectively (see Methods for antibody sources). Anti-Pv12, Pv41 and PvMSP7.1 all showed a clear co-localization with anti-MSP1 (Figure 3A, Figure S2A) suggesting that their orthologous targets are located on the merozoite surface. Anti-PvGAMA and, to a lesser extent, anti-PvCyRPA and anti-PvDBP co-localised with anti-AMA1, suggesting that their orthologous targets are located in apical secretory organelles such as the micronemes (Figure 3B, Figure S3). Anti-ARP appeared to be apically located but did not co-localise with any known markers that we tested (Figure 3B and Figure S3), such that its exact location remains to be determined. To confirm that antibodies against Pv12, Pv41 and PvMSP7.1 were labelling the merozoite surface and not the IMC, which produce similar staining patterns in late schizonts, co-staining with the IMC marker anti-GAP45 was calso arried out in early schizonts, as the IMC and merozoite surface are easier to distinguish earlier in the cell cycle. In all cases there was no co-localisation with anti-GAP45 in early schizonts (Figure S2B), confirming a merozoite surface, not an IMC, location. In all cases the co-localization of the anti-*P*.*vivax* antibody targets as determined by immunolocolisation were identical to those predicted based on their *P. falciparum* homologues.

**Figure 3.**
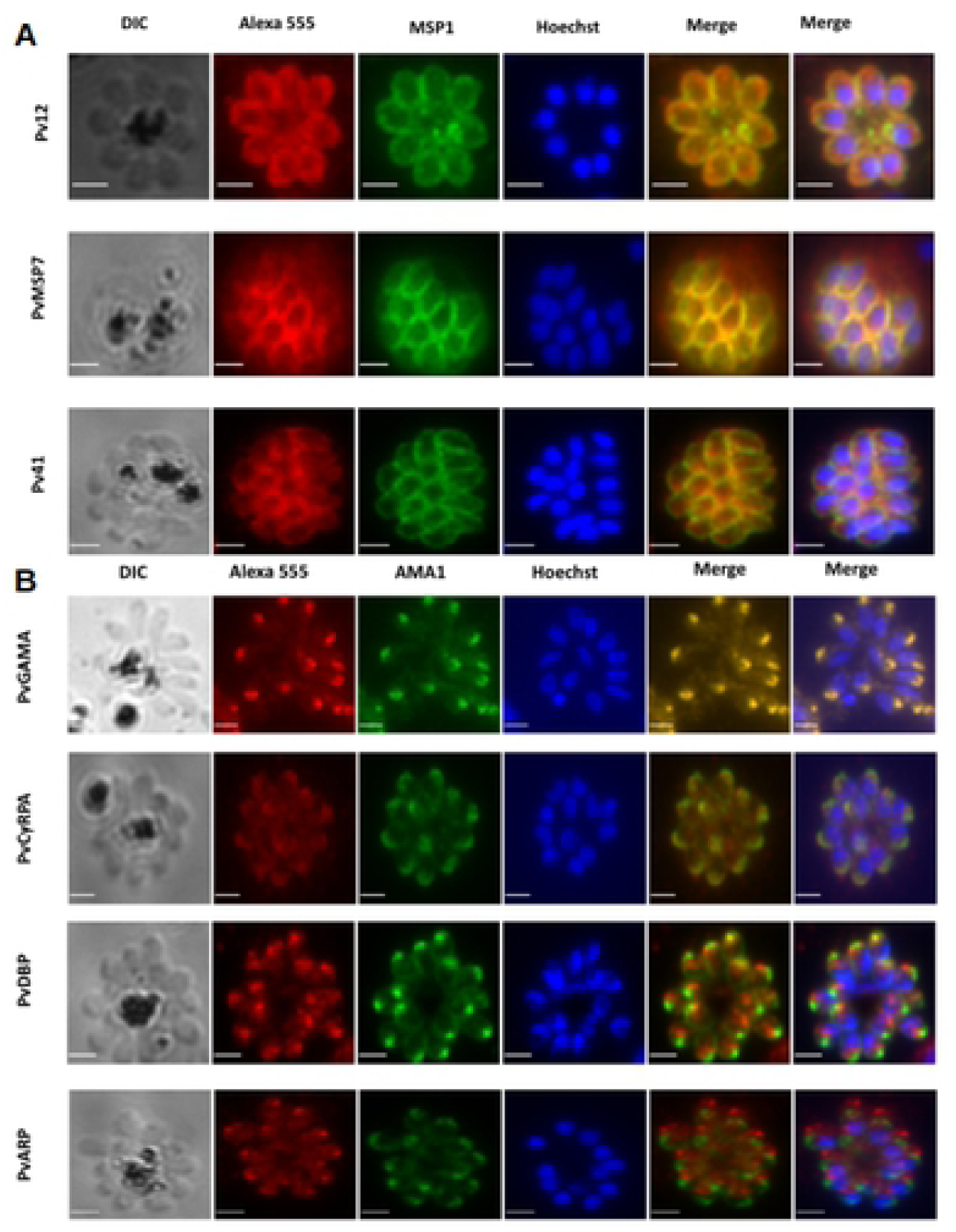
Colocalisation of *P. knowlesi* homologs of *P. vivax* vaccine candidates with antibodies to proteins of known cellular location. *P. knowlesi* proteins were localised using *P. vivax* polyclonal antibodies and compared with A. anti-PkMSP1 or B. anti-PfAMA1 for colocalisation. In both A. and B., Alexa Fluor 555 Goat-anti rabbit and Alexa Fluor 488 Goat-anti rat (Thermo Fisher Scientific) were used as secondary antibodies. Parasite DNA was localised using Hoechst 33342 (Thermo Fisher Scientific). Merge one is an overlay of Alexa 555 and Alexa488, while Merge two is an overlay of Alexa 555, Alexa488 and Hoeschst. Imaging was done using fluorescence microscopy. Scale bar is 2 micrometers.

### Screening anti-*P*.*vivax* antibodies for inhibitory activity in *P. knowlesi* identifies novel invasion-blocking candidates

Having established that anti-*P. vivax* antibodies could be used to specifically detect homologues in *P. knowlesi*, we explored whether the same antibodies could inhibit *P. knowlesi* erythrocyte invasion or intra-erythrocytic development. Serial two-fold dilutions of purified total IgG were prepared starting from 10 mg/ml, and incubated with synchronized ring-stage *P. knowlesi* parasites for 24 hours. Assays were carried out in two biological replicates each with three technical replicates, where invasion and growth inhibition were measured by flow cytometry using Far-red Cell Trace staining of erythrocyte and SYBR green staining of parasite DNA. Invasion was quantified as the percentage of erythrocytes that were both SYBR green and Far-red Cell Trace positive as compared to only control treated erythrocytes, while growth was quantified as the percentage of cells that were only SYBR green positive as compared to control treated erythrocytes. As shown in Figure 4 and Table 2, compared to the positive and negative controls for inhibition (heparin and rabbit IgG respectively), inhibitory activity fell into two broad groups: inhibitory (top panel, anti-Pv12, Pv41, PvGAMA and PvDBP which gave IC_50_ values of 4.17, 11.24, 6.64 and 4.54 mg/ml respectively) and not inhibitory (bottom, anti-PvARP, PvCyRPA, PvMSP7.1, PvMSP3.10). The low level of inhibition observed with anti-PvMSP7.1 and PvMSP3.10 could be due to the low degree of homology between Pv and Pk homologs, and the lack of inhibition observed with the anti-PvMSP3.10 was consistent with the absence of cross-reactivity with PkMSP3 homologues in immunofluorescence assays. Strong inhibition with anti-PvDBP, the only *P. vivax* blood stage vaccine target in the advanced stage of vaccine development, was confirmatory and comparable to other studies (46)⍰. Antibodies to two other targets, Pv12 and PvGAMA, had broadly similar IC_50_s to anti-PvDBP, while antibodies to Pv41, which interacts with Pv12 (32), also had strong inhibition. All three targets were therefore worthy of further investigation.

**Figure 4.**
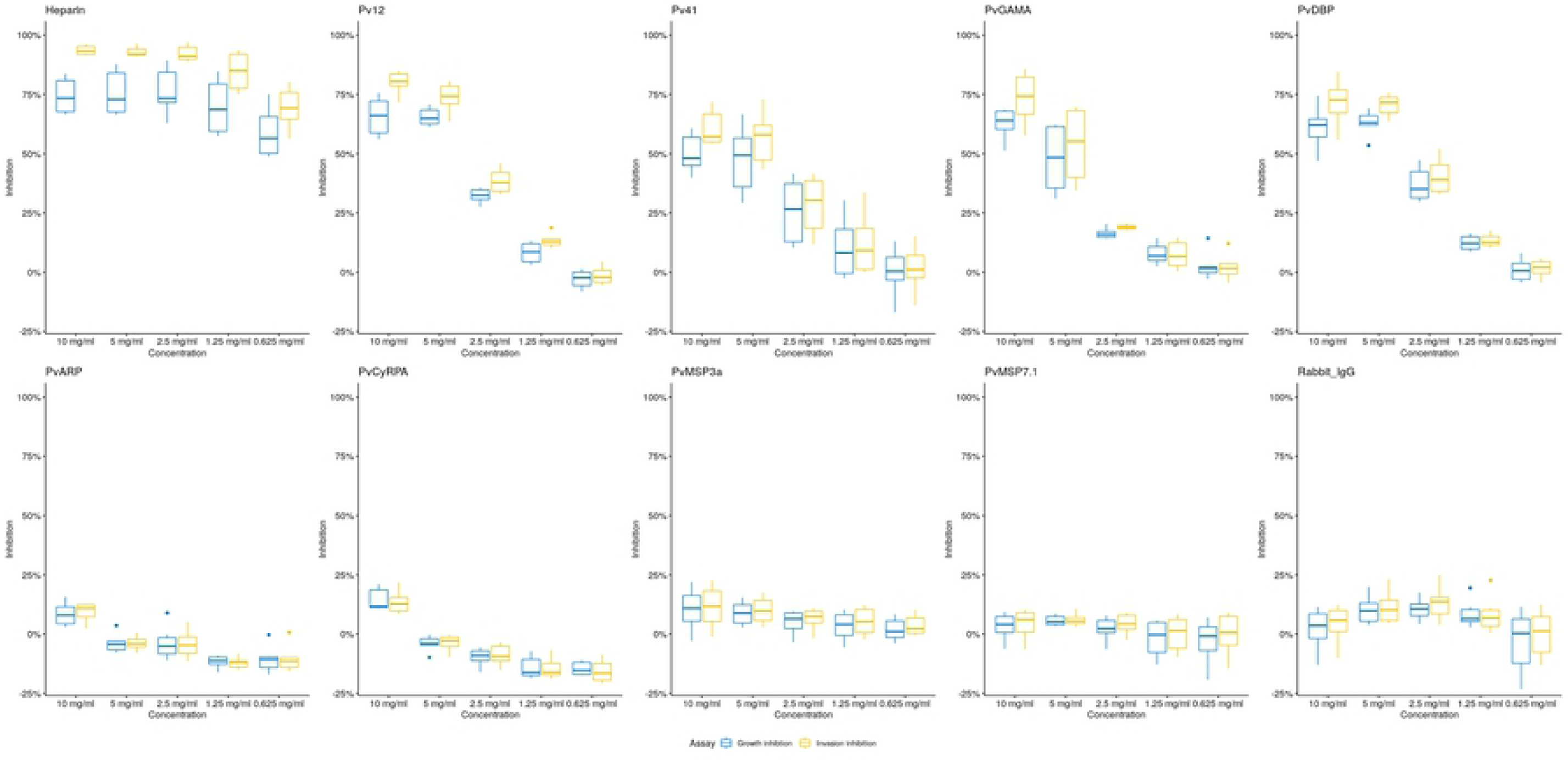
Invasion and growth inhibition of *P. knowlesi* by polyclonal antibodies to *P. vivax* vaccine candidates. Synchronized *P. knowlesi* cultures at ring stage were mixed with Far-red Cell Trace dye treated erythrocytes and DMSO treated erythrocytes. These were then treated with two-fold serial dilutions from 10 mg/ml to 0.625 mg/ml of purified total IgG. Cell numbers were quantified with a flow cytometer using SYBR green after 24 hours. Invasion inhibition was calculated as SYBR green and Far-red Cell Trace positive events while growth was calculated as percentage of SYBR green only positive events. Percentage inhibition of invasion (yellow) and growth (blue) of *P. knowlesi* grown in the presence of antibodies was compared to *P. knowlesi* growth in parallel in the absence of antibodies. Heparin and Rabbit IgG from unimmunized animals were used as positive and negative controls respectively.

### Gene editing in *P. knowlesi* establishes that Pk41 and PkGAMA are not essential for blood-stage growth

Gene essentiality is one potential prioritisation factor in ranking vaccine candidates, as targeting the product of a gene that is absolutely required for parasite development is by definition more likely to yield growth inhibitory activity. Given that antibodies against Pv12, Pv41, and PvGAMA all inhibited *P. knowlesi* growth, we used genome editing to determine whether the orthologous genes in *P. knowlesi* could be knocked out. We also targeted *PkARP* as a positive control, as anti-PvARP antibodies had no inhibitory effects on *P. knowlesi* (Figure 4), while constructs targeting *PkDBPα* were included as a negative control, as this gene has previously been shown to be essential (47)⍰. Gene targeting was carried out using a CRISPR-Cas9 two-vector approach, with one vector containing Cas9 and guide RNA expression cassettes as well as the selection marker, while the other one was a donor template for repair consisting of eGFP flanked by 5’ and 3’ untranslated regions of each respective gene (Figure S4A). Thus, after drug selection based on the selection marker in the guide vector and not in the donor vector, integration of the construct would both eliminate the endogenous gene, and result in eGFP expression. Transfection of *P. knowlesi* was followed by selection with 100 nM pyrimethamine for 6 days to select for Cas9 expression, and cultures were maintained for up to 3 weeks. Transfections were repeated at least twice for each pair of constructs.

Parasites were recovered from all transfections. Genomic DNA was extracted from recovered lines, and used for genotyping to establish whether integration had occurred. Only parasites transfected with *Pk41* and *PkGAMA* knockout constructs gave bands of the size expected if gene deletion had occurred (Figure S4B). Whole genome sequencing analysis confirmed this result, showing no reads mapping to the deleted regions of the wildtype (WT) *P. knowlesi* genome (Figure S5 and S6), indicating that integration of the knock-out construct had occurred. By contrast, genotyping of parasites transfected with *Pk12, PkARP* and *PkDBPα* constructs did not differ from WT cultures. No WT band was amplified for *Pk41* and *PkGAMA* knockout lines, whereas WT parasite controls yielded bands of the expected size (Figure S4B). *Pk41* and *PkGAMA* therefore appear to be non-essential for *P. knowlesi* growth, whereas *Pk12, PkARP* and *PkDBPα* were not able to be disrupted using this approach. For *PkARP* this was unexpected, given that anti-PvARP antibodies had no effect on growth or invasion and the three different guide RNAs used for this gene were effective in targeting this locus in subsequent experiments, shown below.

To confirm that Pk41 and PkGAMA expression was absent in the knockout lines, fluorescence and immunofluorescence assays were performed. Both knockout lines expressed eGFP (Figure 5A and B), while localisation assays with anti-Pv41 and anti-PvGAMA gave no specific signal (Figure 5C), showing only background staining in clear contrast to WT parasites (Figures 3 and 4). This confirms that these parasites were not expressing Pk41 and PkGAMA, and therefore that *Pk41* and *PkGAMA* are redundant for intra-erythrocytic growth, despite the fact that anti-P41 and anti-GAMA antibodies were shown to inhibit parasite growth (Figure 4). Testing the antibodies in growth assays using the knockout strain showed no detectable inhibition, confirming that the antibodies were specific to their immunogens (Figure 6).

**Figure 5.**
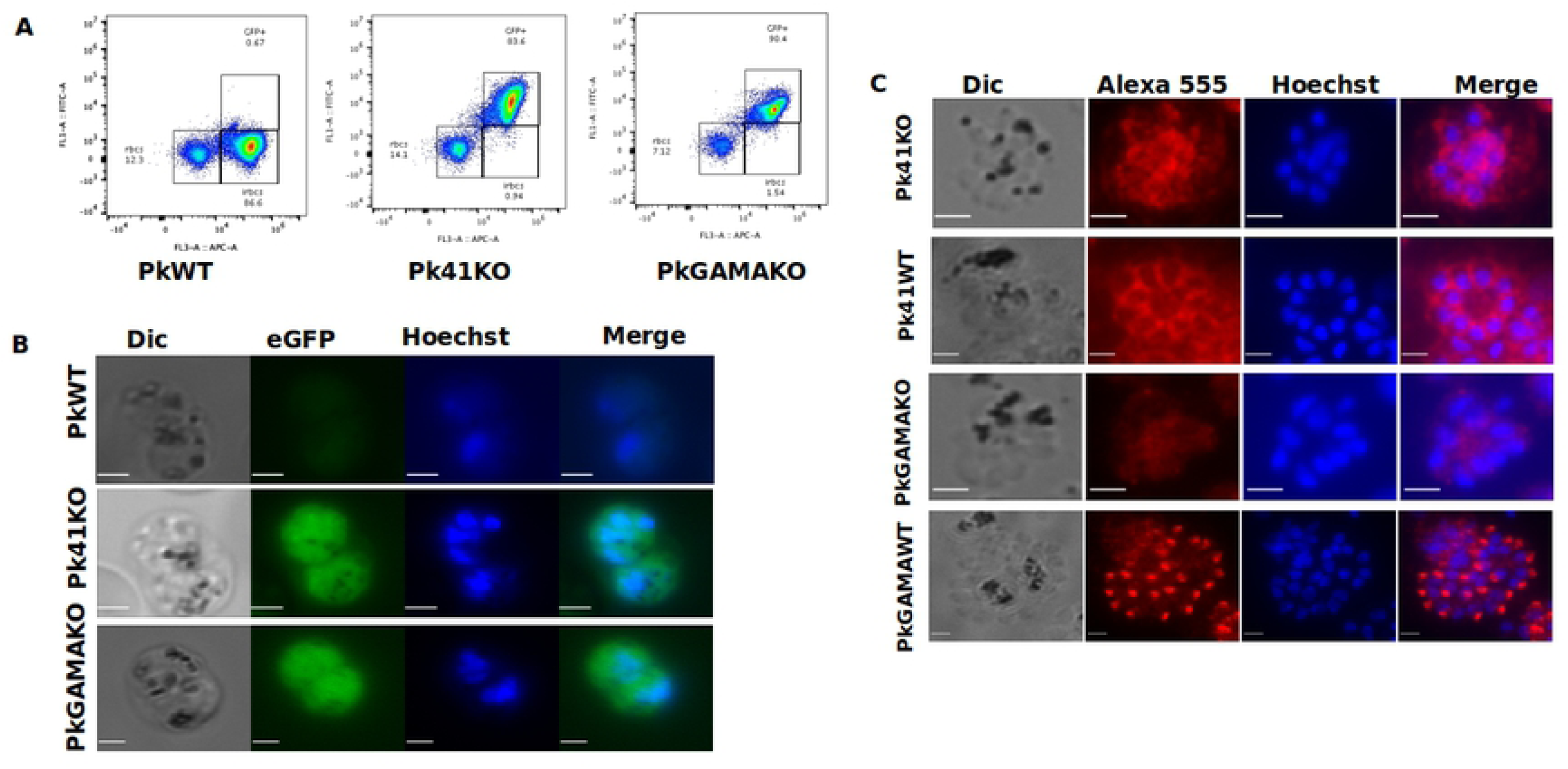
Gene editing can delete PkP41 and PkGAMA in *P. knowlesi*. Pkp41 and PkGAMA were replaced with eGFP to generate Pk41KO and PkGAMAKO strains respectively. A. Flow cytometry to establish eGFP expression in knockout lines. Enriched knock out and wild-type *P. knowlesi* cultures at late stages were treated using SYBR green in 1X PBS and incubated for one hour after which they were quantified by flow cytometry. Events were gated as both SYBR green and GFP negative (Lower left), SYBR green only positive (Lower right) and both SYBR green and GFP positive (upper). B. eGFP expression in knock-out *P. knowlesi* strains as compared to wild-type *P. knowlesi*, imaged using fluorescence microscopy. Parasite nuclei were stained using Hoechst 33342 (Thermo Fisher Scientific). Merge is an overlay of Alexa 555 and Hoeschst. Scale bar is 2 micrometers. C. Proteins in knock out and wild-type *P. knowlesi* strains were localised using *P. vivax* polyclonal antibodies and Alexa Fluor 555 Goat-anti rabbit (Thermo Fisher Scientific) as secondary antibody then imaged using fluorescence microscopy. Localisation in both knock out and wild-type Pv41 and PvGAMA was performed using anti-Pv41 and anti-PvGAMA antibodies respectively. Parasite nuclei were stained with Hoechst 33342 (Thermo Fisher Scientific). Merge is an overlay of Alexa 555 and Hoeschst. Scale bar is 2 micrometers.

**Figure 6.**
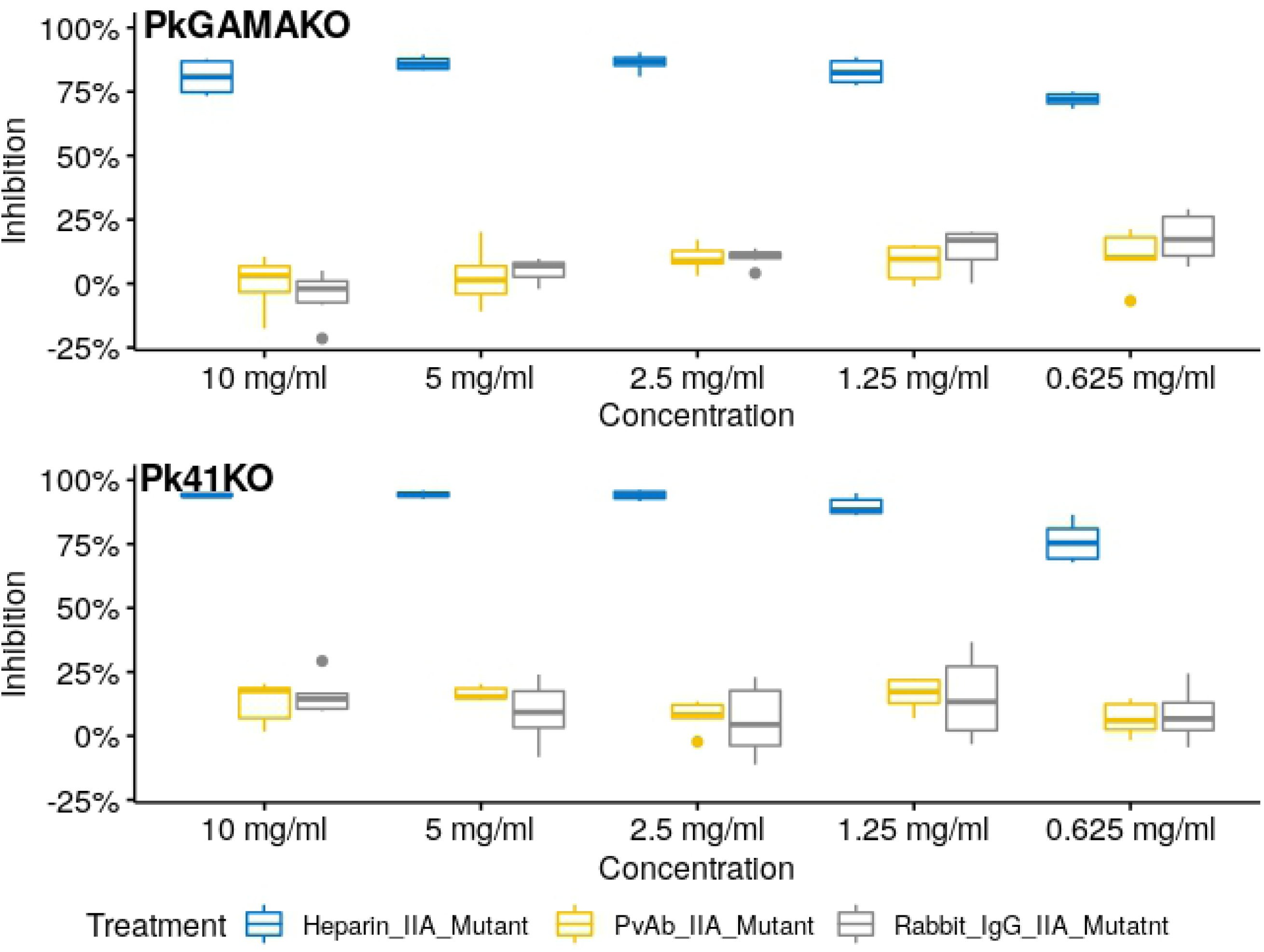
Knockout of *P. knowlesi* genes eliminates inhibition by polyclonal antibodies to homologous *P. vivax* vaccine candidates. Synchronized *P. knowlesi* knockout cultures at ring stage were mixed with Far-red Cell Trace dye treated erythrocytes and DMSO treated erythrocytes. These were then treated with two-fold serial dilutions from 10 mg/ml to 0.625 mg/ml of either Heparin, purified anti-*P. vivax* total IgG or commercial Rabbit IgG. Cell numbers were quantified by a flow cytometryusing SYBR green after 24 hours. Invasion inhibition was calculated as SYBR green and Far-red Cell Trace positive events while growth was calculated as the percentage of SYBR green only positive events. Percentage invasion inhibition of PkGAMAKO and Pk41KO strains treated with Heparin (blue), purified anti-*P. vivax* total IgG (yellow) or commercial Rabbit IgG (grey) compared to untreated *P. knowlesi*. Heparin and commercial Rabbit IgG from unimmunized animals were used as positive and negative controls respectively. Percentage inhibition of *P. knowlesi* under a background of Pk41 and PkGAMA knock out treated with f anti-Pvp41 and anti-PvGAMA antibodies respectively.

### Allele replacement of *P. knowlesi* genes with *P. vivax* orthologues increases the inhibitory effect of anti-*P. vivax* antibodies

A true test of the inhibitory effectiveness of the anti-*P. vivax* antibodies would be in the context of the proteins that they were raised against, but *P. vivax* culture and invasion assays are not available for routine use. To test an alternative approach, we sought to replace *P. knowlesi* target genes with their *P. vivax* orthologues, generating chimeric *P. knowlesi* strains expressing *P. vivax* proteins. Replacement constructs were created in which the *Pv12, Pv41, PvGAMA* and *PvARP* open reading frames were flanked by the 5’ and 3’ UTRs of their *P. knowlesi* counterparts, and these were transfected in combination with the same Cas9/gRNA vectors used in the knockout studies, in order to replace *Pk12, Pk41, PkGAMA* and *PvARP* with *Pv12, Pv41, PvGAMA* and *PvARP* respectively (Figure S7). After selection of transfected parasites with 100 nM pyrimethamine and expansion of the resulting parasites lines, genomic DNA was extracted for genotyping. All lines gave bands of the expected size (Figure S7) indicating that integration of these replacement constructs had occurred at the expected locus, and no WT bands were detected. Whole genome sequencing analysis confirmed that no reads mapped at the targeted region when comparing with Pk reference genome (Figures S8-11).

Localisation assays with anti-Pv12, PvARP, Pv41 and anti-PvGAMA antibodies all gave specific signals in the replacement lines (Figure 7) with anti-Pv12 and Pv41 (Figure 7A and 7C) indicating merozoite surface localisation, while anti-PvGAMA and PvARP (Figure 7B and 7D) appeared as punctate signals, just like signals in wildtype *P. knowlesi* parasites (Figures 2 and 3). These chimeric parasites are therefore viable and able to correctly express and localize Pv12, PvARP, Pv41 and PvGAMA. The chimeric *P. knowlesi* strains had similar growth rates as the WT strains (Figure S12), indicating that the *P. vivax* genes can substitute for the function of their *P. knowlesi* counterparts, emphasizing the phylogenetic relationship between the two parasites. To test whether replacing the *P. knowlesi* genes with their *P. vivax* counterparts increased the inhibitory activity of anti-*P. vivax* antibodies, we tested for growth and invasion inhibition, comparing WT and chimeric replacement lines. In all cases inhibition was increased when using the chimeric lines (Figure 8), indicating that while *P. knowlesi* is a useful model as a first screen for *P. vivax* reverse vaccinology studies, sequence differences between *P. vivax* antigens and their *P. knowlesi* orthologues can lead to underestimation of the inhibitory effect when only wildtype *P. knowlesi* parasites are used.

**Figure 7.**
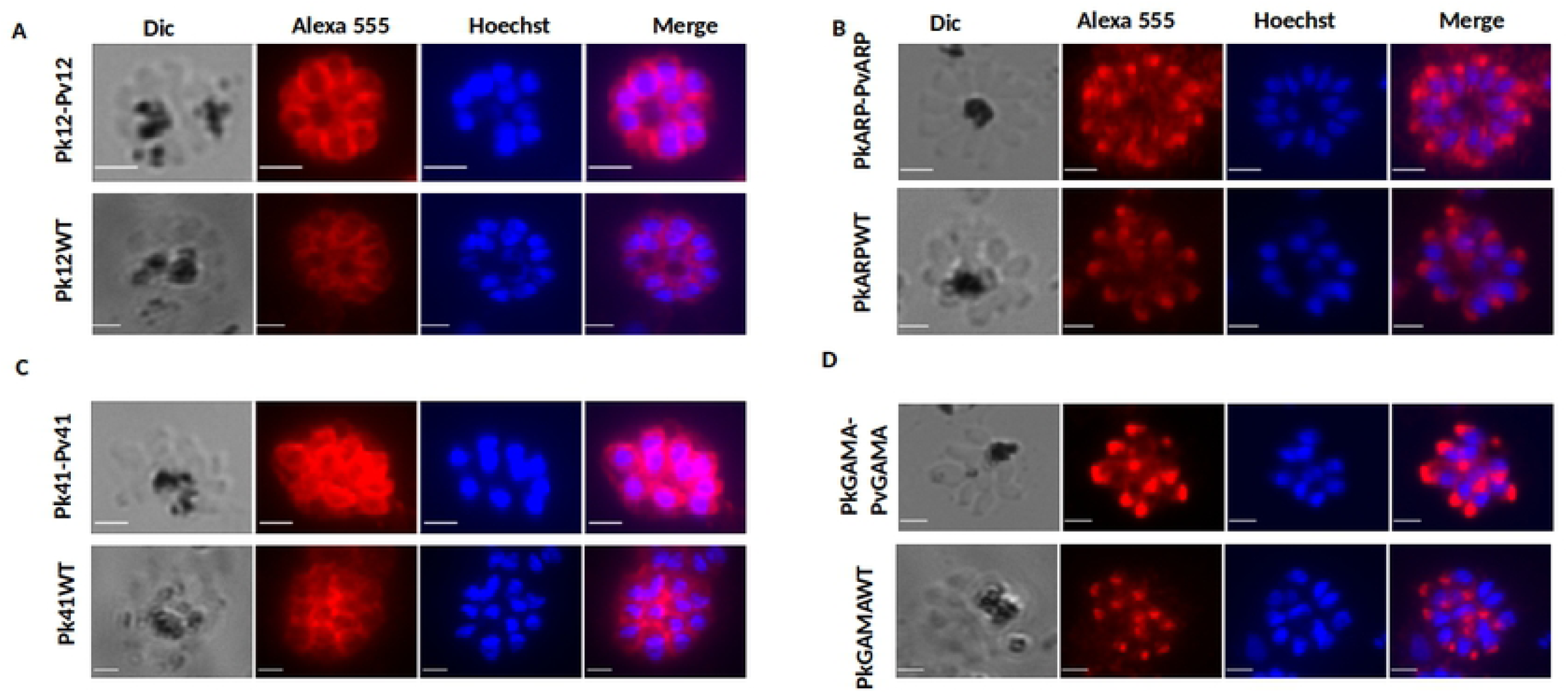
Gene editing to replace *P. knowlesi* target genes with orthologous *P. vivax* candidate genes. Pvp12, PvARP, Ppv41 and PvGAMA gene sequences were used to replace Pkp12, PkARP, Pkp41 and PkGAMA in wild-type *P. knowlesi* to create lines Pk12Rep, PkARPRep, Pk41Rep and PkGAMARep respectively. Proteins in chimeric and wild-type *P. knowlesi* strains were localised using *P. vivax* polyclonal antibodies and Alexa Fluor 555 Goat-anti rabbit (Thermo Fisher Scientific) as secondary antibody and imaged using fluorescence microscopy. Localisation of both chimeric proteins Pv12, PvARP, Pv41 and PvGAMA (Pk12Rep, PkARPRep, Pk41Rep and PkGAMARep respectively) and wild-type proteins Pkp12, PvARP, Pkp41 and PvGAMA (Pk12WT, PkARPWT, Pk41WT and PkGAMAWT respectively) was performed using anti-Pv12, anti-PvARP, anti-Pv41 and anti-PvGAMA antibodies, respectively. Parasite nuclei were stained with Hoechst 33342 (Thermo Fisher Scientific). Merge is an overlay of Alexa 555 and Hoeschst. Scale bar is 2 micrometers.

**Figure 8.**
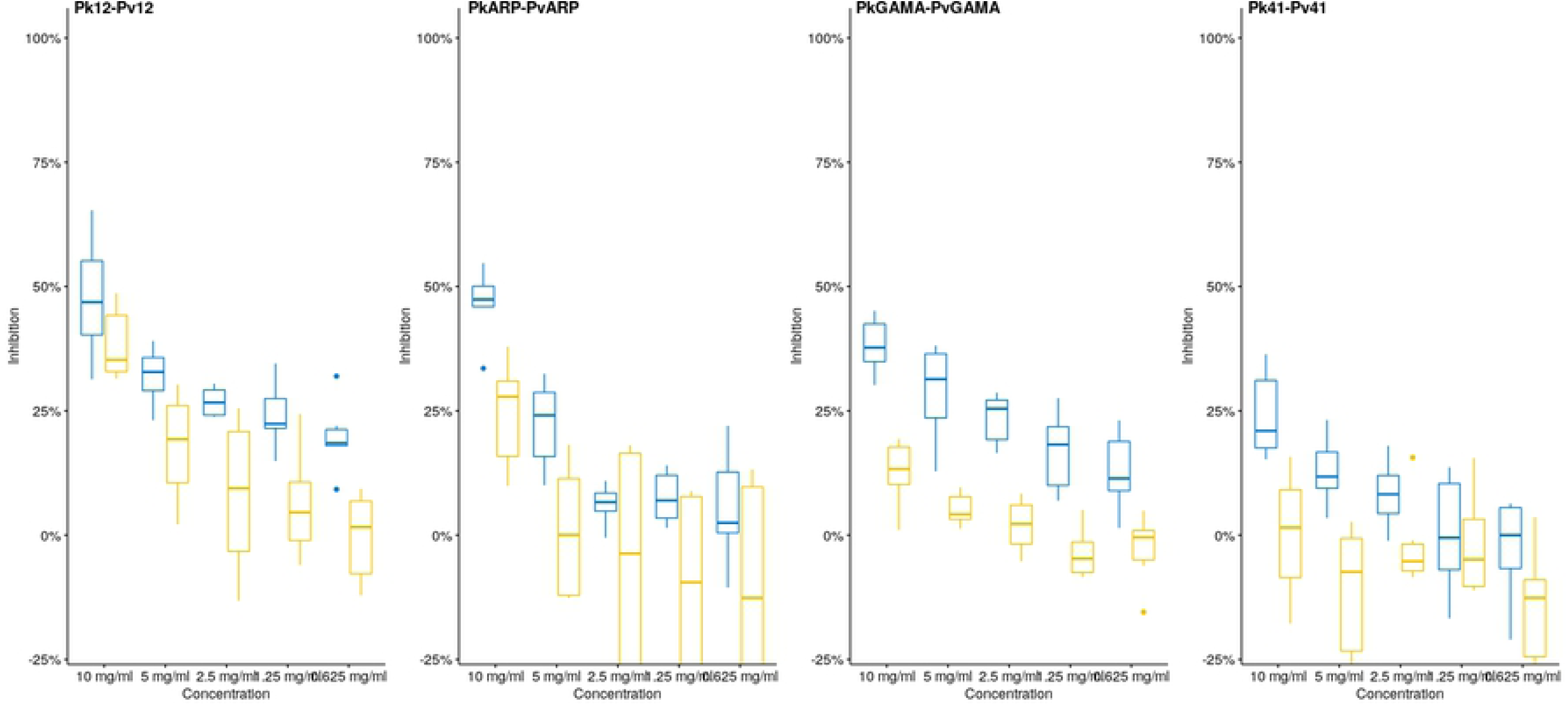
Inhibition of chimeric *P. knowlesi* expressing *P. vivax* proteins by polyclonal antibodies to *P. vivax* vaccine candidates is increased in chimeric *P. knowlesi* strains expressing *P. vivax* proteins. Synchronized wild-type and chimeric *P. knowlesi* strains’ cultures at ring stage were mixed with Far-red Cell Trace dye treated erythrocytes and DMSO treated erythrocytes. These were then treated with two-fold serial dilutions from 10 mg/ml to 0.625 mg/ml of anti-*P. vivax* total IgG. Cell numbers were quantified by flow cytometry using SYBR green after 24 hours. Invasion inhibition was calculated as SYBR green and Far-red Cell Trace positive events while growth was calculated as percentage of SYBR green only positive events. Percentage invasion inhibition of chimeric *P. knowlesi* (blue) expressing Pvp12 (Pkp12Replacement), PvARP (PkARPReplacement), Pvp41 (Pkp41Replacement) and PvGAMA (PkGamaReplacement) treated with anti-Pvp12, anti-PvARP, anti-Pvp41 and anti-PvGAMA antibodies respectively compared to *P. knowlesi* WT (yellow).

## DISCUSSION

To date only a limited number of *Plasmodium vivax* blood stage vaccine candidates have been investigated (reviewed in (6,7,48,49)). This is in large part because it is currently not possible to continuously culture *P. vivax* blood stages *in vitro*, which rules out many biological assays. We have explored whether *P. knowlesi*, which has a close phylogenetic relationship with *P. vivax* (39,40) and has been adapted to *in vitro* culture in human erythrocytes (41,50), could be used to screen for *P. vivax* blood-stage vaccine candidates, as it can for drug-resistance candidates (51). Such an approach has proven viable to explore the most advanced *P. vivax* blood-stage vaccine candidate, PvDBP (46). In this case we sought to apply the *P. knowlesi* model to systematically screen for new blood-stage antigens, using a panel of polyclonal antibodies generated against candidates from a previously published library of *P. vivax* schizont expressed proteins (32). A similar approach has recently been applied to a panel of *P. vivax* blood stage targets, although functional testing did not include knockout and gene replacement strategies (52). We focused on seven *P. vivax* targets: two merozoite surface proteins (PvMSP7.1 and PvMSP3.10); two 6-Cysteine domain proteins (Pv12 and Pv41) and three proteins not belonging to other families (PvGAMA, PvCyRPA and PvARP), with PvDBP included as a positive control.

There was a broad correlation between the ability of anti-*P. vivax* antibodies to specifically recognise their *P. knowlesi* orthologues in immunoblot and immunofluorescence assays and the percent identity within the *P. vivax/P. knowlesi* antigen pairs. However, it is worth noting that a single dominant protein band was identified in 5/8 cases, and clear intracellular localisations defined in 7/8 cases, despite levels of identity as low as 50%, suggesting the system has broader utility than homology levels alone might indicate. This high level of cross-reactivity may be in part be due to our strategy of raising polyclonal antibodies against full-length protein ectodomains, whereas many antigen studies focus only on smaller sub-domains, which limits the chances that cross-reactive responses will be generated. There is also some evidence that the eukaryotic expression system we use increases the likelihood of generating antibodies against folded, functional domains (43), which are more likely to be of utility in assays such as immunofluorescence or growth inhibition, where conformation-dependent epitopes are more important. Using these antibodies in growth inhibition assays revealed robust dose-dependent inhibition of *P. knowlesi* growth by anti-Pv12, Pv41 and PvGAMA antibodies, in some cases on a similar level to anti-PvDBP.

Pv12 and Pv41 are members of a family of 6-cysteine domain proteins, other members of which are under investigation as transmission-blocking vaccine targets in *P. falciparum* (53–55). The *P. falciparum* orthologues of Pv12 and Pv41 form a heterodimer and are localised on the merozoite surface (56). We have previously shown that Pv12 and Pv41 are also able to heterodimerize (32), and here we show that their *P. knowlesi* orthologues also colocolise to the merozoite surface, suggesting key elements of the function of these two proteins are conserved across *Plasmodium* species. However, there are also elements that are different. Immunoepidemiology studies show that antibody responses to Pv12 and Pv41 are commonly induced by exposure to *P. vivax* infection (32,57–59), and have been associated with protection against severe *P. vivax* malaria, in keeping with the inhibitory activity of anti-Pv12 and Pv41 antibodies shown here. By contrast, in *P. falciparum*, anti-Pf12 and Pf41 antibodies have no inhibitory effects on parasite growth *in vitro*, and the genes can be deleted, suggesting a level of functional redundancy in this species (56). Previous immunofluorescence studies of Pv12 in *P. vivax* suggest that it localises to the rhoptries, rather than merozoite surface as in our experiments (60)⍰, although whether this apparent difference is due to differences in the stage of the parasite during erythrocytic schizogony used in the assays is not known.

We localised PkARP to the apical region of the merozoite, suggesting a possible location in the rhoptries like the PfARP homologue in *P. falciparum* (60), but in contrast to previous studies suggesting a merozoite surface location in *P. vivax* and *P. knowlesi* (52,61); again, experimental differences in the stage of parasites used might explain the different observations. PkGAMA also localised to apical organelles, replicating previous observations of a micronemal location in *P. vivax (29)* and *P. falciparum* (62,63). We were unable to inhibit *P. knowlesi* growth using anti-PvARP antibodies, contradicting what has been shown in *P. knowlesi* (64) and *P. falciparum* (60). However, when we replaced PkARP with PvARP, there was a reversal of the activity of anti-PvARP, suggesting that PkARP may lack key inhibitory epitopes recognised by our polyclonals, which were raised against PvARP. Anti-PvGAMA had invasion and growth inhibitory effects on *P. knowlesi* in both WT and chimeric lines, replicating the observations of anti-PfGAMA effects on *P. falciparum* (63), suggesting a conserved role of this protein in invasion across species. Immunoepidemiology studies also show that antibody responses to PvARP and PvGAMA are commonly induced by exposure to *P. vivax* infection (14,32,57–58,65), and have been associated with protection against severe *P. vivax* malaria.

How do these relatively new candidates compare to the much more well-studied vaccine candidate PvDBP? Clearly PvDBP has been the subject of decades of work, meaning that there are multiple lines of evidence supporting its candidacy. The limitations of this target are also well known, specifically the challenge of strain-specific antibody responses, which may be able to be overcome with epitope engineering (28,66–68). One question in weighing up the candidacy of any vaccine antigen is whether the gene that encodes it is essential for parasite growth, as targeting a non-essential gene would seem likely to select for parasites that do not rely on the gene product, and so are able to escape the vaccine.

According to the current model, PvDBP is essential for invasion, as *P. vivax* primarily invades reticulocytes via the interaction between PvDBP and its host receptor, Duffy Antigen Receptor for Chemokine (DARC) (17–20). However, it has now clearly been demonstrated that *P. vivax* is also able to infect individuals who are Duffy negative, lacking DARC expression on their red blood cell surface (21,22). This could indicate that *P. vivax* is able to utilize other ligands for invasion such as PvRBP2b (28) and erythrocyte binding protein 2 (ebp2) (30), although it is also possible that PvDBP is still involved in the invasion of Duffy negative cells (26). It is also worth noting that the genetic essentiality of *PvDBP* for parasite growth has never been able to be directly tested, as *P. vivax* cannot be cultured and therefore cannot be genetically manipulated. Studies in *P. knowlesi*, which has three homologues of *PvDBP*, suggest that at least one is required for invasion of human erythrocytes (47), but whether this is true of *PvDBP* remains to be proven unequivocally.

In the case of the four candidates identified by initial screening with wildtype *P. knowlesi*, there are clear 1:1 orthologues between *P. vivax* and *P. knowlesi*, providing an even stronger rationale than that of Pv/PkDBP to use *P. knowlesi* genetic tools to explore candidates. We utilised the fact that *P. knowlesi* can be readily genetically manipulated (46) to explore whether Pv12, Pv41, PvGAMA and PvARP were essential for parasite growth. *Pk41* and *PkGAMA* could be experimentally deleted, while *Pk12* and *PkARP* could not, even though both anti-PvP41 and PvP12 antibodies had invasion and growth inhibitory effects on WT *P. knowlesi*. The fact that *Pk41* and *PkGAMA* could be deleted without any apparent effect on growth, whereas antibodies that recognise them inhibit growth, seems contradictory. This could be explained if the antibodies raised against Pv41 and PvGAMA recognized multiple targets in *P. knowlesi*, but these antibodies had no inhibitory activity when incubated with the relevant knockout strains, nor could they detect any protein in immunofluorescence assays in these lines. This strongly suggests that the antibodies are specific, and rules out off-target explanations for the antibody inhibition data. An alternative explanation is that the process of genetic deletion, which takes some weeks to recover modified parasites, provides an opportunity for the parasites to adapt to the loss of a specific gene, for example by up-regulating the expression of other genes. By contrast, growth inhibition assays occur in a single cycle, which the parasites may find it harder to adapt to. Vaccine-induced antibodies would arguably operate in a similar manner, suggesting that while essentiality might be one element used to prioritise new targets, it should definitely not be the only one. Despite this, it would seem reasonable to argue that Pv12 and PvARP should have a higher priority for follow-up than Pv41 and PvGAMA.

This study clearly highlights several advantages of the *P. knowlesi* system as a model for testing *P. vivax* blood-stage antigens, as has been suggested in previous drug (51) and vaccine (46,52) studies. A key one is accessibility -access to *ex vivo P. vivax* samples is limited and samples are precious, whereas we were readily able to perform multiple *in vitro* assays using *P. knowlesi*. A second is also the genetic accessibility of the system, where gene deletions and allele replacements, while not precisely routine, are certainly readily achievable. There are however always limitations in using one species as a model for another, and this study reveals some of these. The study relies on antibodies that were generated against *P. vivax* proteins being able to cross-react with their *P. knowlesi* orthologues, and while in almost all cases this proved possible, there was some correlation between the strength of cross-reactivity and the % identity between antigen pairs, meaning the *P. knowlesi* model will almost certainly be more useful for some antigens than others. The genetic tractability of *P. knowlesi* offers a potential solution to this problem, as we have shown, by allowing the replacement of endogenous *P. knowlesi* genes with their *P. vivax* orthologues, implying that antibodies can be tested against the precise sequence they were generated against. This approach relies on the ability of a *P. vivax* gene to substitute for *P. knowlesi* gene function, which may not always be the case, but in all four instances tested here, as well as the Pv/PkDBP swaps carried out by others (46), this has not proven a problem. Ultimately however, no model system is perfect, even *in vitro* culture of *P. vivax* itself, which after all is only a model for *in vivo* growth. It would be extremely useful to the *P. vivax* vaccine field to carry out a head-to-head comparison across all the four currently available functional models - wildtype *P. knowlesi*, genetically modified *P. knowlesi* with allele modifications to insert *P. vivax* genes, *P. cynomologi* and *P. vivax ex vivo* assays. The targets identified here, Pv12, Pv41, PvARP and PvGAMA, along with PvDBP, present a perfect opportunity to carry out such a test.

To conclude, using both antibody and genetic approaches, we exploited the phylogenetic relationship between *P. knowlesi* and *P. vivax* to explore blood-stage *P. vivax* vaccine targets. The data suggests a hierarchy of possible targets, with Pv12 and PvARP being the highest priority as they are genetically essential and can be targeted with inhibitory antibodies, Pv41 and PvGAMA in a second tier as they can be inhibited with antibodies but also genetically deleted, while PvMSP7.1, PvMSP3.10 and PvCyRPA would seem to have the lowest priority. We have demonstrated that antibodies against *P. vivax* vaccine targets are able to recognise proteins in *P. knowlesi* as well as inhibit its growth and invasion, and that *P. knowlesi* has many advantages as a rapid and accessible system to screen *P. vivax* blood stage targets.

These advantages need to be balanced against the limitations described above, and it is always possible that lack of inhibition and in some case lack of localisation in *P. knowlesi* using anti-*P. vivax* antibodies is due to the lower level of similarity between *P. vivax* and *P. knowlesi* orthologues, or indeed differences in biology between these species. Until a robust *P. vivax* culture system is established, which despite extensive effort by multiple teams (34–38)⍰ does not seem likely soon, it will be advisable to use multiple models to screen for candidates, and be clear and upfront about the limitations inherent in all of them.

## MATERIALS AND METHODS

### *In vitro* parasite culture

*Plasmodium knowlesi* parasites were maintained as described in (41). Briefly, *P. knowlesi* strain A1-H.1 was propagated in human O^+^ erythrocytes (UK NHS Blood and Transplant), in RPMI 1640 supplemented with Albumax (Thermo Fisher Scientific), L-Glutamine (Thermo Fisher Scientific), Horse serum (Thermo Fisher Scientific), Gentamicin (Thermo Fisher Scientific). The cultures were kept at 2% hematocrit, gassed using a mixture of 5% CO_2_, 5% O_2_ and 90% Nitrogen, while being monitored three times per week by counting parasitemia using light microscopy with media change or splitting as appropriate.

### Synchronization and enrichment of *Plasmodium knowlesi*

Synchronization was performed by enriching late stage parasites using Histodenz (Sigma Aldrich) as described in (41). Briefly, parasites were resuspended in 5ml complete media and layered on top of 5 ml of 55% Histodenz in complete culture media in a 15 ml tube (Greiner). The mixture was then centrifuged for 3 minutes at room temperature, 1500 g, acceleration 3 and brake 1, resulting in late stage parasites becoming enriched at the interface. For inhibition assays, these parasites were returned to culture, and assays set up after reinvasion had occurred in the subsequent cycle. For protein extracts, immunofluorescence assays and transfections, this was repeated over three consecutive cell cycles to create very tightly synchronized parasites, with schizont samples from a fourth cycle of Histodenz purification used for subsequent analysis.

### Antibody production and purification

Rabbits were immunized with 1mg of his-tagged *P. vivax* full-length ectodomain, expressed in HEK239E cells as previously described (32), and purified by nickel affinity chromatography. Immunisations were carried out using Freunds complete/incomplete adjuvant by Cambridge Research Biochemicals. Total IgG was purified using Protein G gravitrap kit (GE healthcare). Eluted total IgG was concentrated by centrifugation at 4°C for 30 mins using 100000MWCO vivaspin (Sartorius). The concentrate was then dialyzed using a dialysis tube (Millipore) overnight at 4°C with 1 litre of RPMI 1640 (Thermo Fisher Scientific), before repeating concentration if necessary. Total IgG concentration was measured bynanodrop (NanoDrop).

### Protein extraction and Western blotting

To generate protein extracts, schizont stage parasites enriched from 5-10 mL of culture at 5-10% parasitemia were treated with 0.15% saponin (Sigma Aldrich) and protease inhibitor (Cat. No. 5892970001, Sigma Aldrich) at 1X for 1min on ice to release parasites from their host cell. After pelleting and two rounds of washing in ice cold 1X PBS (Sigma Aldrich) with protease inhibitor 1X, parasites were treated with 1 µl of DNAse I (Thermoscientific) for 30mins at 37°C., before mixing 1:1 with Laemmli sample buffer 2X and incubating for 30mins-1h at 37°C to gently denaturate the sample. The pellet was then frozen down at -80°C until needed. Samples were diluted 0, 1:5 or 1:10 in Laemmli, and 5µl of the diluted samples were loaded onto a 4-12% bis-tris NuPage gel (Thermo Fisher Scientific) and run at 200V for 50min in MOPS buffer (Thermo Fisher Scientific). For electrophoresis, 1 µg/µl of samples was mixed with 2.5 µl NuPage LDS sample buffer (4X) (Thermo Fisher Scientific), 1 µl NuPage Reducing agent (10X) (Thermo Fisher Scientific) and deionized water to 6.5 µl. The mixture was then incubated at 72°C for 10 mins while shaking at 300 RPM. 10 µl of the sample was then resolved on a 4-12% bis-trisNuPage gel (Thermo Fisher Scientific) with 1X MOPS SDS gel buffer (Thermo Fisher Scientific) at 200 V for 50 minutes.

Western blot transfer was carried out in wet conditions at 30V for 60min, and membrane blocked overnight while shaking at 4°C in 5% milk (Marvel) containing 0.077% sodium azide (Sigma Aldrich). Primary antibodies were diluted in 5% milk/PBS/0.1% TWEEN 20 at concentrations as follows: anti-PvGAMA 1:2400; Pvp12, PvMsp7.1 and PvMSP3.10 1:400; Pvp41, PvARP and PvCyRPA 1:800; PvDBP 1:1600. Primary incubation was carried out overnight at 4°C. Blots were then washed three times, each 10mins, in PBST (1X PBS and 0.1% Tween), before incubating with secondary anti-Rabbit HRP (Abcam) at 1:20000 dilution in 5% milk/PBS/0.1% for 45 mins at room temperature. Blots were washed again three times, each 10mins, in PBST before developing with ECL prime Western Blotting detection reagent, (GE Healthcare). Expected molecular weight of the *P. vivax* candidate proteins and their orthologous proteins in *P. knowlesi* was predicted using Protein Molecular weight calculator (69) based on the amino acid sequences of the respective protein sequences from PlasmoDB.

### Immunofluorescence assays

Cells were synchronized, harvested from culture and enriched using Histodenz as described above based on (41). These were then washed with 1X PBS (Sigma Aldrich) for 5 mins and fixed with 4% paraformaldehyde (Agar scientific)/ 0.0075% glutaraldehyde (Sigma Aldrich) in 1X PBS for 30 minutes at room temperature, followed up with washing in 1x PBS while shaking for 5 mins. Thin smears of fixed cells were made on Poly-L-Slides (Sigma Aldrich) and stored in -80°C freezer until needed. For processing, slides were incubated briefly at room temperature, permeabilized with 0.1% Triton X-100 (Sigma Aldrich) in 1X PBS for 30 mins at room temperature then washed once with 1x PBS while shaking for 5 mins. Blocking was carried out overnight at 4°C in a humidified dark chamber using Blocking Aid Solution (Thermo Fisher Scientific). Primary antibody diluted in Blocking Aid Solution (Anti-PvGAMA 1:1200; Pvp12, PvMsp7.1 and PvMSP3.10 1:200; PvARP 1:400; PvCyRPA and PvDBP 1:800; Pvp41 1:200, PkMSP1-19 1:2000; PfAMA1 1:1000) was then added and incubated overnight in a humidified dark chamber at 4°C followed by washing three times for 5 mins in 1X PBS while shaking. Secondary antibody diluted in Blocking Aid Solution (Alexa Fluor 555 Goat-anti rabbit (Thermo Fisher Scientific) (1:500) for anti-*P. vivax* rabbit polyclonals and Alexa Fluor 488 Goat-anti rat (Thermo Fisher Scientific) (1:500) for PkMSP1-19 and PfAMA1) was then added and incubated 1 hour in a humidified dark chamber at room temperature followed by washing three times for 5 mins in 1X PBS while shaking. Hoechst 33342 (Thermo Fisher Scientific), for nucleus staining, was diluted 1:3000 in 1x PBS (Sigma Aldrich) then added and incubated for 10 mins in a humidified dark chamber at room temperature with subsequently washing three times for 5 mins in 1X PBS while shaking. The cells were later mounted with Pro-Long Gold mounting solution (Thermo Fisher Scientific), covered with cover-glass (VWR), left to cure for 24 hours in dark and dry chamber at room temperature and eventually sealed with slide sealer (Biotium), before imaging on a Leica DMi8.

### Invasion and growth inhibition assays

Two milli-litres of O+ erythrocytes at 2% hematocrit in incomplete-culture media were labelled using 2 µl of a stock of 1 mM Far-red Cell Trace (Thermo Fisher Scientific); control unstained erythrocytes were incubated with 2 µl of Dimethyl-sulphoxide (DMSO; Sigma Aldrich). After 2 hours of incubation at 37°C while shaking, labelled erythrocytes were washed twice using complete media, then the cells were resuspended in complete media to 2% hematocrit in 2 ml final volume. Labelled erythrocytes were mixed with synchronized rings, generated by enriching schizonts as described above then returning them to culture until reinvasion had occurred. The labelled erythrocyte-parasite mix was incubated with serial dilutions of anti-*P. vivax* antibodies, with all dilutions made using incomplete medium. Incubations were carried out in 96-well plates, with each well containing 20 µl infected erythrocytes, 20 µl stained erythrocytes, Xµl of diluted total IgG (Xµl because the antibodies were in varying stock concentrations therefore requiring different volumes to be added to get the same final concentrating), and 2.2 µl of a mixture of serum, hypoxanthine and gentamicin (at a ratio of 2, 0.18 and 0.009 respectively). Control wells contained 40 µl of erythrocytes only, or 20 µl of infected erythrocytes/20 µl of unstained erythrocytes, or 40 µl of stained erythrocytes only, or 20 µl of infected erythrocytes and 20 µl of stained erythrocytes, to control for gating in the flow cytometry. Triplicates of each combination were incubated in a 96 well plate for 24 hours in a gassed chamber. To quantify parasite invasion or growth, samples were centrifuged for 3 mins at 450g (acceleration 9, brake 3) at room temperature, supernatant was removed and samples were labelled with SYBR green I nucleic acid dye (Thermo Fisher Scientific) for 1 hour at 37°C while shaking at 52 rpm. After two washes with 100 µl 1 X PBS (Sigma Aldrich), samples were resuspended in 100 µl of 1 X PBS (Sigma Aldrich) and parasites quantified using FACS (Cytoflex, Becton and coulter) as previously described (70). Data was analyzed using FlowJo (FlowJo)then using Excel (Microsoft office), invasion was calculated as the percentage of erythrocytes that were both SYBR green and Far-red Cell Trace positive as compared to only DMSO treated erythrocytes while growth was calculated as the percentage of cells that were only SYBR green positive as compared to only DMSO treated erythrocytes. The results were then plotted using the following R packages; ggplot2 (71)⍰, ggpubr (72), cowplot (73), magrittr (74), readxl (75) and dplyr (76) in R-Studio (R-Studio Inc). IC_50_ was determined using Probit/logit regression using Excel (Microsoft office).

### Genetic modification

#### Gene repair construct design and assembly

Constructs, guide RNAs and primers (supplementary table S1,2,4,5,6) were designed with (77) and (78). Synthetic DNA codon optimization was performed using gblocks® Gene Fragments (IDT) (PvGAMA_regions1 & 3, PvARP_region1) and GeneArt Gene Synthesis (Thermo Fischer Scientific) (Pv12, Pv41, PvARP_region3). PvGAMA_regions2 and PvARP_regions3 were amplified from expression constructs previously generated by (32)⍰. Other fragments were amplified from *P. knowlesi* genomic DNA, purified from saponin-lysed *P. knowlesi* infected erythrocytes using DNA blood kit (Qiagen) according to the manufacturer’s protocol. Gene editing donor vectors were assembled in PUC19 using Gibson assembly according to the manufacturer’s protocol (NEB), using PCR products amplified using KAPA HiFi HotStart ReadyMixPCR Kit (KAPABiosystems) and purified using gel isolation kits (Macherey and Nagel). Primers (Supplementary Tables S1 and S2) and PCR programs (Supplementary Table S3: KAPA2M for all of constructs except KAP121M for final amplification of *Pkgama* replacement insert) are listed in the Supplementary Material.

#### Assembly of Cas9/gRNA vectors

The cloning vector (TGL96) was digested using BtgZI (NEB), purified using a gel purification kit (Macherey and Nagel), and treated with shrimp alkaline phosphatase (NEB) to dephosphorylate vector ends. Guide RNAs (Supplementary Table S5) were reconstituted by mixing 1 µl of 100 µM stocks of the forward and reverse strands for each guide with 1 µl of 10x ligation buffer (NEB), 0.5 µl T4 polynucleotide kinase (NEB) and 65 µl nuclease free PCR water. Annealing was carried out by incubating at 37°C for 30 min, then increasing to 94°C for 5 min before cooling at 25°C at a ramp speed of 5°C per-min. Annealed primers were then diluted to 1 µl in 200 µl and ligated (NEB) to the digestedand dephosphorylated Cas9 vector.

Vectors were transformed into chemically competent *E. coli* according to the manufacturer’s protocol (NEB), and grown overnight. Resulting colonies were screened by colony PCR using GoTaq Green PCR master mix (Promega), with 1 µl of Pk5’ UTR forward and Pk3’UTR reverse primers for each respective construct. Positive colonies were expanded and DNA purified using a miniprep purification kit (Macherey and Nagel) and sequenced to confirm construct integrity (GATC/Eurofins). Sequencing data was analyzed using Benchling (77) and Seqman Pro (DNA star Navigator); positive constructs were expanded and purified for transfection using a maxiprep purification kit (Macherey and Nagel).

### Transfection

Transfection was performed largely as described in (41). Late stage *P. knowlesi* parasites were enriched using Histodenz as described above. In each transfection cuvette (Lonza), 10 µl of schizonts was mixed with 100 µl of P3 solution (Lonza) containing 30 µg each of the relevant donor and guide vectors. Transfections were carried out using program FP158 (Amaxa Nucleofector, Lonza), and the contents were then immediately transferred into a 2 ml sterile eppendorf tube containing 500 µl of complete culture media mixed with 190 µl uninfected erythrocytes. The transfection mix was incubated at 37°C while shaking at 800 rpm in a thermomixer for 30mins, before being transferred into a 6 well plate, gassed and incubated at 37°C for one parasite life cycle. After this selection was applied with 100 nM pyrimethamine (Santa Cruz Biotechnology Inc). For three days, the cultures were monitored by smearing and selection done by changing the media and replacing with fresh media containing 100 nM pyrimethamine (Santa Cruz Biotechnology Inc). On day 4 post transfection the cultures were diluted 1/3 in 5ml fresh media containing 100 nM pyrimethamine and 100 µl erythrocytes. The cultures were then maintained and monitored after every 2nd cycle by smearing/parasitemia counting, with media changed or cultures split as appropriate. Samples where parasites re-appeared were expanded in a total volume of 50 ml with erythrocytes at 2% hematocrit until parasitemia was greater than 5%. DNA was then isolated using a DNA Blood kit (Qiagen) and analysed using gene-specific primers (Supplementary Table S6) and PCR program KAPA18C (Supplementary Table S3). Cultures that contained only modified parasites were phenotyped without cloning, while those that genotyping showed had both modified and WT genotyping bands were cloned by limiting dilution and plaque cloning in flat-bottomed 96-well plates. Wells containing single plaques were identified using an EVOS microscope (4x objective, transmitted light), expanded, and DNA isolated and genotyped as described above as well as whole genome sequenced WGS analysis was then performed on the Welcome Sanger Cluster using bowtie2 (79), samtools (80). Visualisation was perfomedusing Integrative Genomics Viewer (81–83) as described in Pevner (84)⍰.

### Ethics

Human O+ erythrocytes were purchased from NHS Blood and Transplant, Cambridge, UK, and all samples were anonymised. The work complied with all relevant ethical regulations for work with human participants. The use of erythrocytes from human donors for *P. falciparum* culture and binding studies was approved by the NHS Cambridgeshire 4 Research Ethics Committee (REC reference 15/EE/0253) and the Wellcome Sanger Institute Human Materials and Data Management Committee.

## ACKNOWLEDGEMENTS

We wish to acknowledge the following for their various contributions: Ellen Knuepfer for the kind donation of Rat anti-PkMSP1-19 serum. Rob Moon and Franziska Mohring for the kind donation of Pk CRISPR-Cas9, guide and donor vectors, and advice on genetic modification, Mehdi Ghorbal for advice on construct design, cloning and genetic manipulation experiments and Allan Muhwezi for advice on cell culture.

Funding was provided by the National Institutes of Health (R01AI137154), European Union (MultiViVax 773073) and the Wellcome Trust (206194/Z/17/Z). The funders had no role in study design, data collection and analysis, decision to publish or preparation of the manuscript.

The views expressed in this article are those of the authors and do not necessarily represent the views of the National Heart, Lung and Blood Institute, the National Institutes of Health, the United States Department of Health and Human Services, or any other government entity.

## Supporting information

**S1 Supplementary figure 1. Preimmune antibody controls for immunolocalization**. Localisation of PkGAMA, PkDBP, PkARP, PkCyRPA, PkMSP7.1, Pk41 and Pk12 using pre-immune serum from rabbits immunized with PvGAMA, PvDBP, PvARP, PvCyRPA, PvMSP7.1, Pv41 and Pv12, respectively. Scale bar is 2 micrometers.

**S2 Supplementary figure 2. Colocalization of *P. knowlesi* proteins using polyclonal anti-*P. vivax* antibodies to *P. vivax* vaccine candidates with antibodies to proteins of known cellular location**. Colocalization of Pk12, PkMSP7.1, Pk41 with A. PkMSP1-19 and B. PkGap45 using antibodies to Pv12, PvMSP7.1, Pv41, PkMSP1 and PfGap45, respectively. Scale bar is 2 micrometers.

**S3 Supplementary figure 3. Colocalization of *P. knowlesi* proteins using polyclonal anti-*P. vivax* antibodies to *P. vivax* vaccine candidates with antibodies to proteins of known cellular location**. Colocalization of PkGAMA, PkCyRPA, PkDBP, PkARP, Pk12, PkMSP7.1, Pk41 with **A)** PkMSP-1-19 and **B)** PkAMA1 using anti-bodies to PvGAMA, PvCyRPA, PvDBP, PvARP, Pv12, PvMSP7.1, Pv41, PkMSP1 and PfAMA1, respectively. Scale bar is 2 micrometers.

**S4 Supplementary figure 4. Gene editing strategy to knock out candidate genes**. A) General strategy used to knock out *Pk12, Pkarp, Pkdbpalpha, Pk41* and *Pkgama*. Plasmids used are Cas9/gRNA vector and donor vector containing eGFP (GFP) flanked with 5’ and 3’ untranslated region (UTR) for each respective *P. knowlesi* gene (PkCDS). Primer pairs used for genotyping are P1&P2 and P3&P4, to test for wildtype, P1&P5 and P6&P4 to test for integration of the knockout construct. B) Genotyping of Pk41KO and PkGAMAKO with above primer pairs as compared to WT. On the right side are the obtained molecular weight in kilobase pairs (kb).

**S5 Supplementary figure 5. Whole genome sequencing of Pk41 knockout strain**. Alignment of p41 knockout strains to *P. knowlesi* reference genome and the WT strain from which they were generated.

**S6 Supplementary figure 6. Whole genome sequencing of PkGAMA knockout strain**. Alignment of GAMA knockout strains to *P. knowlesi* reference genome and the WT strain from which they were generated.

**S7 Supplementary figure 7. Gene editing strategy to replace *P. knowlesi* target genes with orthologous *P. vivax* candidate genes**. A) General strategy used to replace *pk12, pkarp, pk41* and *pkgama* with *pv12, pvarp, pv41* and *pvgama*, respectively. Plasmids used are Cas9/gRNA vector and donor vector containing the *P. vivax* coding sequence (PvCDS) flanked with 5’ and 3’ untranslated region (UTR) for each respective *P. knowlesi* gene (PkCDS). Primer pairs used for genotyping are P1&P2 and P3&P4, to test for WT. P1&P5 and P6&P4 to test for integration of the replacement construct. B) Genotyping of pk12, pkarp, pk41 and pkgama allele replacement (Rep) using the above primer pairs as compared to WT. On the right side are the obtained molecular weight in kilobase pairs (kb).

**S8 Supplementary figure 8. Whole genome sequencing of Pk41-Pv41 replacement strain**. Alignment of p41 allele replacement strains to *P. knowlesi* reference genome and the WT strain from which they were generated.

**S9 Supplementary figure 9. Whole genome sequencing of PkGAMA-PvGAMA replacement strain**. Alignment of GAMA allele replacement strains to *P. knowlesi* i reference genome and the WT strain from which they were generated.

**S10 Supplementary figure 10. Whole genome sequencing of PkARP-PvARP replacement strain**. Alignment of ARP allele replacement strains to *P. knowlesi* i reference genome and the WT strain from which they were generated.

**S11 Supplementary figure 11. Whole genome sequencing of Pk12-Pv12 replacement strain**. Alignment of p12 allele replacement strains to *P. knowlesi* i reference genome and the WT strain from which they were generated.

**S12 Supplementary figure 12. Comparative Growth rate assay between wildtype *P. knowlesi* and genetically edited *P. knowlesi strains*. WT** (P. knowlesi WT), Gamako (PkGama knock-out clone), Gamarep (PkGama replacement clone), p41ko (Pkp41 knock out clone), p41rep (Pkp41 replacement clone), P12rep (Pkp12 replacement clone), ARPrep (PkARP replacement clone).

